# Selective Functional Connectivity between Ocular Dominance Columns in the Primary Visual Cortex

**DOI:** 10.1101/2024.05.22.595395

**Authors:** Iman Aganj, Shahin Nasr

## Abstract

The primary visual cortex (V1) in humans and many animals is comprised of fine-scale neuronal ensembles that respond preferentially to the stimulation of one eye over the other, also known as the ocular dominance columns (ODCs). Despite its importance in shaping our perception, to date, the nature of the functional interactions between ODCs has remained poorly understood. In this work, we aimed to improve our understanding of the interaction mechanisms between fine-scale neuronal structures distributed within V1. To that end, we applied high-resolution functional MRI to study mechanisms of functional connectivity between ODCs. Using this technique, we quantified the level of functional connectivity between ODCs as a function of the ocular preference of ODCs, showing that *alike* ODCs are functionally more connected compared to *unalike* ones. Through these experiments, we aspired to contribute to filling the gap in our knowledge of the functional connectivity of ODCs in humans as compared to animals.

## 1 Introduction

Resting-state functional connectivity (rs-FC) has been extensively used to study the large-scale functional organization and networks of the human brain. However, the use of rs-FC at the mesoscale level – especially to study brain circuitry – has been more limited [1-4]. This is mainly due to the technical challenges in visualizing mesoscale neuronal ensembles and measuring their rs-FC reliably across scan sessions, e.g. in the case of ocular dominance columns (ODCs) [5-7] that are small compared to the spatial resolution of conventional functional neuroimaging techniques.

Visual processing relies on functional interactions between fine-scale neuronal clusters distributed across the visual cortex. Within the primary visual cortex (V1; one of the most studied areas in the visual system), ODCs are considered the building blocks of visual processing, with interconnections [8-10] that have an important role in shaping our perception [11-16]. Anatomical studies in animals have shown that ODCs are tightly interconnected through *selective* horizontal connections [17-21]: ODCs that share ocular preference (i.e., with *alike* preference) exhibit stronger connections to each other than those with *unalike* ocular preference [8-10]. Despite direct relevance to a wide range of perceptual impairments (e.g. amblyopia [22-24]), however, our knowledge of rs-FC between ODCs and its role in controlling perception in *humans* is limited. Providing evidence for homologous connections in the human V1 via measuring the extent and selectivity of functional connections between ODCs would fill the gap in our understanding of the functional organization of ODCs in humans as compared with animals.

New data acquisition and processing technologies have recently enabled the study of mesoscale rs-FC in humans. In this work, by focusing on the human V1 organization, we assess the capabilities of our high-resolution functional MRI (fMRI) techniques in revealing rs-FC and its selectivity between ODCs. Specifically, we test the hypothesis (based on animal models [16, 17, 25-28]) that rs-FC between ODC pairs varies with respect to their ocular preference, showing that ODCs with alike ocular preference are more strongly interconnected than those with unalike preference.

We present our methods in Section 2, and provide our results and discuss them in Section 3.

## 2 Methods

### 2.1 Data Acquisition

#### Participants

11 subjects (3 females), aged 25-45 years old, participated in this study. All participants had radiologically intact brains and no history of neuropsychological disorders. One subject was excluded from the study because their ODCs were not detected reliably. All experimental procedures conformed to the guidelines of the National Institutes of Health and were approved by the Massachusetts General Hospital protocols. Written informed consent was obtained from all participants prior to all experiments.

#### Procedure

For ODC localization, participants were scanned 2-3 times in an ultra-high field 7T scanner (whole-body system, Siemens Healthcare, Erlangen, Germany) for the functional experiments. They were then scanned in a separate session to measure their rs-FC. All participants were also scanned in a 3T scanner (Tim Trio, Siemens Healthcare) for structural imaging.

#### ODC Mapping

We stimulated the participant’s left and right eyes in different blocks (i.e., block-design; 24 s per block). The stimuli were sparse (5%) moving random red (50% of blocks) and green (the rest of blocks) dots (0.09°×0.09°; 56 cd/m^2^), presented against a black background. During the fMRI experiments, stimuli were presented via an LCD projector with a 1024×768 (H×W) pixel resolution and 60 Hz refresh rate onto a rear-projection screen, viewed through a mirror mounted on the receive coil array. MATLAB (MathWorks, Natick, MA, USA) and the Psychophysics Toolbox [29, 30] were used to control stimulus presentation.

Participants viewed the stimuli through custom-made anaglyph spectacles (with red and green filters) mounted to the head coil. During the blocks, dots were oscillating horizontally (−0.22° to 0.22°; 0.3 Hz). Stimuli extended 20°×26° in the visual field. Each experimental run began and ended with 12 s of uniform black. The sequence of blocks was pseudo-randomized across runs (14 blocks per run) and each subject participated in 12 runs. Filter laterality (i.e., red-left vs. red-right) was counter-balanced between sessions and across participants. The participants were instructed to look at a centrally presented fixation object (radius = 0.15°) and to do either a shape-change for the fixation target (circle-to-square or vice versa) during the ocular dominance activity measurements or a random dot-detection during the retinotopic mapping.

#### Retinotopic Mapping

For all participants, the border of V1 was defined retinotopically [31]. Stimuli were based on a flashing radial checkerboard, presented against a gray background within retinotopically limited apertures, including wedge-shaped apertures radially centered along the horizontal and vertical meridians (polar angle = 30°). These stimuli were presented to participants in different blocks pseudo-randomly sequenced across runs (24 s per block and 8 blocks per run, for at least 4 runs).

#### Imaging

Functional experiments (see above) were conducted in a 7T whole-body Siemens scanner equipped with SC72 body gradients (70 mT/m maximum gradient strength and 200 T/m/s maximum slew rate) using a custom-built 32-channel helmet receive coil array and a birdcage volume transmit coil. Voxel dimensions were nominally 1.0 mm. We used single-shot gradient-echo EPI to acquire functional images with the following protocol parameter values: TR=3000 ms, TE=28 ms, flip angle=78°, matrix=192×192, BW=1184 Hz/pix, echo-spacing=1 ms, ⅞ phase partial Fourier, FOV=192×192 mm, 44 oblique-coronal slices, acceleration factor *R*=4 with GRAPPA reconstruction and FLEET-ACS data [32] with 10° flip angle. The field of view included occipital cortical areas V1–V4.

Resting-state data were acquired during 4, 3, and 1 runs for 4, 5, and 1 subjects, respectively, with each run containing 128 time points. To test the reproducibility, two subjects were rescanned on a different day.

Structural (anatomical) data were acquired in a 3T Siemens TimTrio whole-body scanner, with the standard vendor-supplied 32-channel head coil array, using a 3D T1-weighted MPRAGE sequence with protocol parameter values: TR=2530 ms, TE=3.39 ms, TI=1100 ms, flip angle=7°, BW=200 Hz/pix, echo spacing=8.2 ms, voxel size=1.0×1.0×1.33 mm^3^, and FOV=256×256×170 mm^3^.

### 2.2 Data Analysis

Functional and anatomical MRI data were pre-processed and analyzed using FreeSurfer and FS-FAST (version 7.11; http://surfer.nmr.mgh.harvard.edu) [33] and in-house MATLAB code.

#### Structural Analysis

For each participant, inflated and flattened cortical surfaces were reconstructed based on the high-resolution anatomical data [33]. During this process, the standard pial surface was generated as the border between the gray matter (GM) and the surrounding cerebrospinal fluid (CSF). The white matter (WM) surface was also generated as the WM-GM interface. To enable intra-cortical smoothing (see below), we also generated a family of 9 intermediated equidistant surfaces, spaced at intervals of 10% of the cortical thickness, between the WM and pial surfaces. To improve the co-registration of functional and structural scans, all surfaces were upsampled [34].

#### Functional Analysis

The collected functional data were first upsampled (to 0.5 mm isotropic) and then corrected for motion artifacts. For each participant, functional data from each run were rigidly (6 DoF) aligned relative to their own structural scan using rigid Boundary-Based Registration [35]. This procedure enabled us to compare data collected for each participant across multiple scan sessions.

To retain the spatial resolution, no tangential spatial smoothing was applied to the imaging data acquired at 7T. Rather, we used the method of radial (intracortical) smoothing [36] – i.e., perpendicular to the cortex and within the cortical columns. For deep cortical depths, the extent of this radial smoothing was limited to WM-GM interface and the adjacent 2 surfaces right above it (see above) – i.e., the bottom 30% of the GM thickness. For the superficial cortical depths, the extent of this procedure was limited to GM-CSF interface and the adjacent 2 surfaces right below it. For the middle cortical layers, the extent of this procedure was limited to the three middle reconstructed cortical surfaces.

A standard hemodynamic model based on a gamma function was fitted to the fMRI signal to estimate the amplitude of the blood-oxygen-level-dependent (BOLD) response. For each individual participant, the average BOLD response maps were calculated for each condition. Finally, voxel-wise statistical tests were conducted by computing contrasts based on a univariate general linear model, and the resulting significance maps were projected onto the participant’s anatomical volumes and reconstructed cortical surfaces.

#### Resting-State Data Analysis

To measure rs-FC, partial correlation coefficients of the resting-state data were computed for pairs of vertices within the V1 cortical surface mesh of each brain hemisphere, reconstructed for each subject based on their own structural data (see above). Head motion and WM signals were used as nuisance regressors.

The selectivity of functional connections between ODCs was inferred by comparing the level of correlation between vertices with alike vs. unalike ocular preference. This was done both in the entire V1 (of each hemisphere) as well as in separate ODC distance quantiles. In the latter case, given the known GE-BOLD point spread function in deep cortical layers [37, 38], we only included the rs-FC values between ODCs at least 3 mm apart from each other.

Next, we tested the reproducibility of the rs-FC selectivity by comparing it in two sessions of each of the two subjects for which a second session (on a different day) had been acquired. We further made use of these two-day rs-FC data to assess the ability of machine-learning approaches to predict the ODC map from the rs-FC patterns in V1, using one rs-FC session for training and the other session for prediction. To that end, we employed machine learning methods implemented in MATLAB’s Regression Learner app (such as variations of the linear regression, tree, support vector machine, efficient linear, ensemble, Gaussian process regression, multilayer neural network, and kernel methods).

## 3 Results and Discussion

### ODC Mapping

We measured the evoked ocular dominance activity for all participants in the deep cortical depth level across visual areas by subtracting the response of the non-dominant eye from the response of the dominant eye. The evoked activity is shown in **Fig. 1** for a representative participant. Consistent with post-mortem anatomical studies in humans [39] and non-human primates [40-42] with normal vision, the cortical topography of the evoked ocular dominance response was organized into mostly parallel interdigitated stripes. These fine-scale stripes – formed by ODCs with alike preference – were similarly detected across cortical depths, reflecting the columnar organization of V1 ODCs [40]. In both hemispheres, these stripes were predominantly limited to the regions of V1 (radius < 10°), representing the central retinotopic visual field that was stimulated during the scans, whereas they were absent in other visual areas. This pattern was consistently observed in all participants in each hemisphere.

**Fig. 1.**
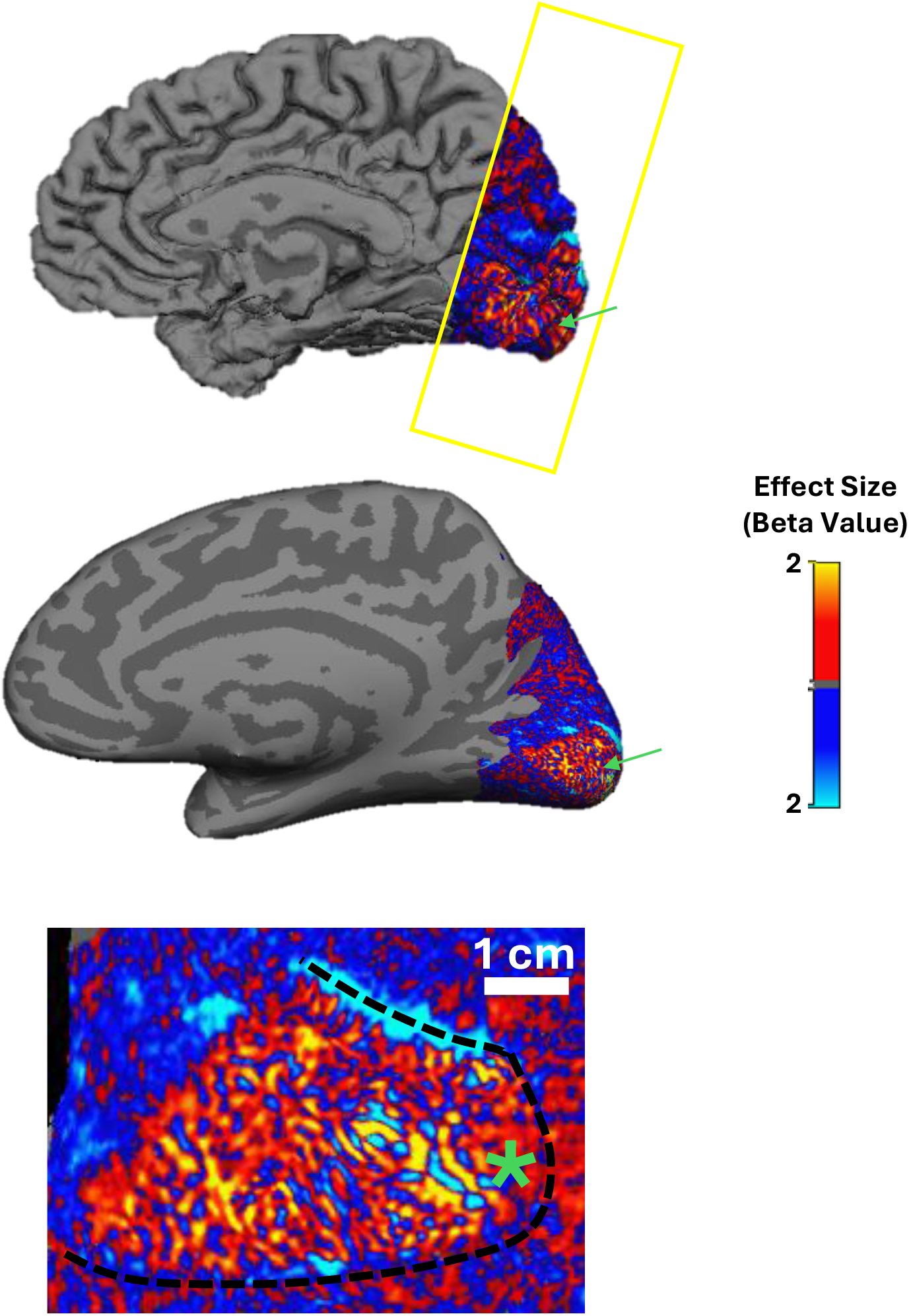
ODC maps made by contrasting the response to left vs. right eye stimulation, overlaid on the pial (top), inflated (middle), and flattened (bottom) representations of a subject’s cortex. The green arrow and asterisk indicatethe foveal direction. The black dashed line shows the borders of V1, defined retinotopically.

### Selectivity of rs-FC

The level of rs-FC between ODCs with alike (same) vs. unalike (different) ocular preference is shown in **Fig. 2** for both hemispheres of all 10 subjects. Paired *t*-tests across subjects showed mean rs-FC to be significantly higher between vertices with alike than unalike ocular preference in the left (*p*=0.007) and right (*p*=0.004) hemispheres.

**Fig. 2.**
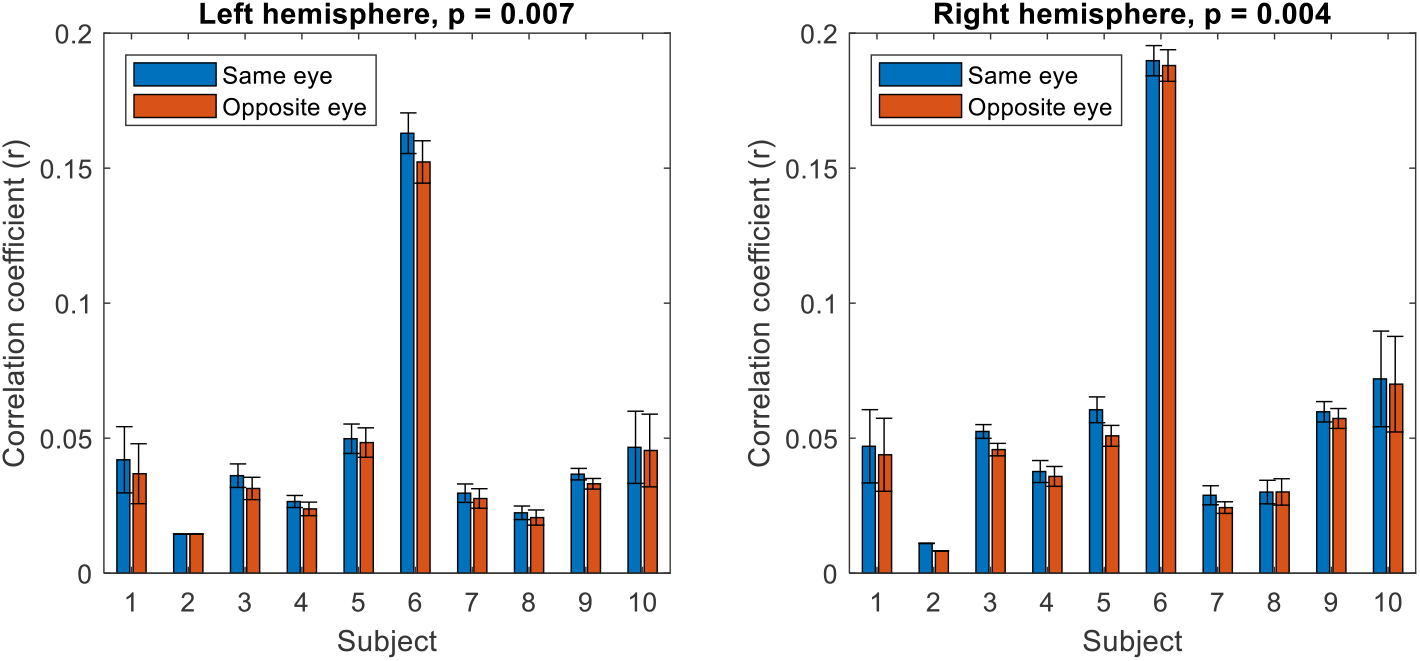
The level of rs-FC between ODCs with alike (same eye) vs. unalike (opposite eye) ocular preference, averaged across runs (error bars: standard error of the mean) for each subject and hemisphere. Paired *t*-test across subjects in each hemisphere revealed mean rs-FC to be significantly higher between vertices with alike rather than unalike ocular preference.

Mean rs-FC for ODC pairs at different distance brackets, averaged across subjects and hemispheres, is shown in **Fig. 3**. The overall level of rs-FC between ODCs decreased as their distance from each other increased. Importantly, the significantly stronger correlation values between ODCs with alike rather than unalike ocular preference were observed even at longer distances. These results support the main hypothesis that, as in animals, human ODCs with alike ocular preference exhibit stronger rs-FC to each other, even across a long distance.

**Fig. 3.**
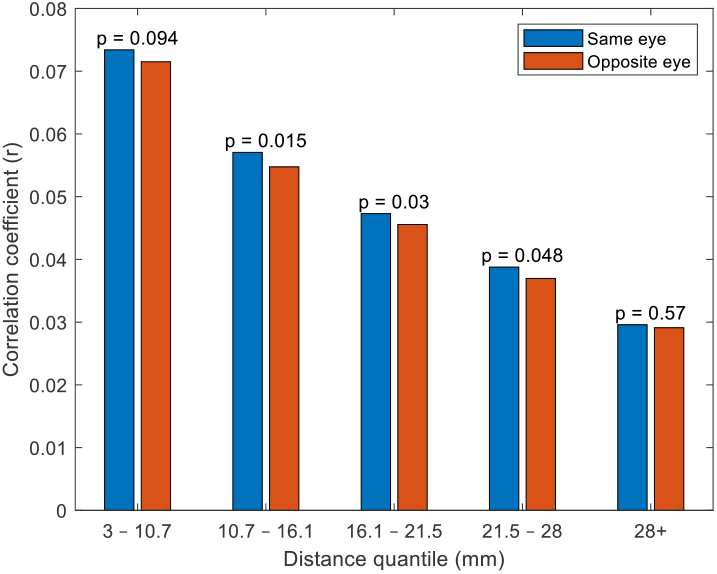
The level of rs-FC between ODCs with same vs. different ocular preference, averaged across hemispheres and subjects, for different distance brackets. Paired *t*-tests are across subjects (hemispheres averaged).

### Reproducibility

The rs-FC correlation values, computed *vertex*-wise, are smaller than commonly reported *region*-wise rs-FC, where within-region signal averaging increases the signal-to-noise ratio. Despite the low levels, the selective rs-FC between ODCs was consistent across different runs within each scan session, as reflected in the small standard error of their means (**Fig. 2**). To test the reproducibility of this phenomenon from day to day, two subjects were re-scanned on a different day. As **Fig. 4** shows, in both subjects, despite the slight change in the level of rs-FC between ODCs across sessions, the selective trend of rs-FC (i.e., stronger for alike than unalike ODCs) remained consistent.

**Fig. 4.**
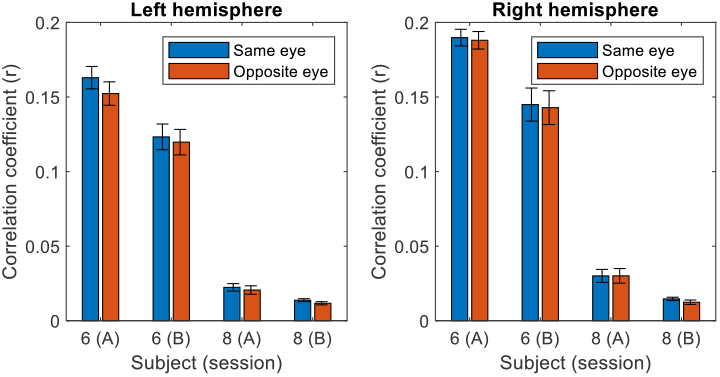
Reproducibility of rs-FC selectivity between ODCs across two scan sessions for two subjects. See **Fig. 2** for details.

We further tested the informativeness of rs-FC in predicting the ODC map. We trained machine-learning models using the rs-FC of the V1 measured in one day, and then tested it with rs-FC acquired on a different day. This test was separately applied to the hemispheres of two subjects for whom rs-FC was collected on two different days; hence, 2 subjects × 2 hemispheres × 2 train/test directions = 8 experiments. The best-performing method was the ensemble of learners (with bootstrap aggregation), resulting in Pearson correlations between the predicted and ground-truth ODC beta maps that were positive in all 8 experiments (two-sided sign test *p*=0.008), with a mean (± standard error) of *r*=0.15±0.02.

### Conclusions

We used the well-studied functional organization of ODCs to test the capabilities of rs-FC in human subjects to reveal the mesoscale functional organization of the human visual cortex. Our results, when combined with the findings based on anatomical [8-10] and rs-FC measurements in animals [16, 17, 25-28], can help to fill the gap in our knowledge of rs-FC between ODCs in humans as compared to animals. Such knowledge may be beneficial to future clinical studies by providing new potential biomarkers for clinicians who desire to monitor the impact of various visual disorders on the fine-scale functional organization of the visual cortex.

## Acknowledgments

Support for this research was provided by the National Institutes of Health, specifically the National Eye Institute (R01EY030434), as well as the National Institute on Aging (RF1AG068261) and the Michael J. Fox Foundation for Parkinson’s Research (MJFF-021226).

## References

1. Kennedy, B., Bex, P., Hunter, D., Nasr, S.: Two fine-scale channels for encoding motion and stereopsis within the human magnocellular stream. Prog. Neurobiol. 220, 102374 (2023)

2. Tootell, R.B., Nasr, S.: Scotopic Vision Is Selectively Processed in Thick-Type Columns in Human Extrastriate Cortex. Cereb. Cortex 31, 1163–1181 (2021)

3. Tootell, R.B., Nasr, S.: Columnar Segregation of Magnocellular and Parvocellular Streams in Human Extrastriate Cortex. J. Neurosci. 37, 8014–8032 (2017)

4. Nasr, S., Polimeni, J.R., Tootell, R.B.: Interdigitated Color- and Disparity-Selective Columns within Human Visual Cortical Areas V2 and V3. J. Neurosci. 36, 1841–1857 (2016)

5. Nasr, S., Skerswetat, J., Gaier, E.D., Malladi, S.N., Kennedy, B., Tootell, R.B., Bex, P., Hunter, D.G.: Using high-resolution functional MRI to differentiate impacts of strabismic and anisometropic amblyopia on evoked ocular dominance activity in humans. bioRxiv 2024.2002. 2011.579855 (2024)

6. Yacoub, E., Shmuel, A., Logothetis, N., Ugurbil, K.: Robust detection of ocular dominance columns in humans using Hahn Spin Echo BOLD functional MRI at 7 Tesla. Neuroimage 37, 1161–1177 (2007)

7. Cheng, K., Waggoner, R.A., Tanaka, K.: Human ocular dominance columns as revealed by high-field functional magnetic resonance imaging. Neuron 32, 359–374 (2001)

8. Tychsen, L., Ming-Fong Wong, A., Burkhalter, A.: Paucity of horizontal connections for binocular vision in V1 of naturally strabismic macaques: Cytochrome oxidase compartment specificity. J. Comp. Neurol. 474, 261–275 (2004)

9. Yoshioka, T., Blasdel, G.G., Levitt, J.B., Lund, J.S.: Relation between patterns of intrinsic lateral connectivity, ocular dominance, and cytochrome oxidase-reactive regions in macaque monkey striate cortex. Cereb. Cortex 6, 297–310 (1996)

10. Malach, R., Amir, Y., Harel, M., Grinvald, A.: Relationship between intrinsic connections and functional architecture revealed by optical imaging and in vivo targeted biocytin injections in primate striate cortex. Proceedings of the National Academy of Sciences 90, 10469–10473 (1993)

11. Polat, U., Sagi, D.: Spatial interactions in human vision: from near to far via experience-dependent cascades of connections. Proceedings of the National Academy of Sciences 91, 1206–1209 (1994)

12. Polat, U., Norcia, A.M.: Neurophysiological evidence for contrast dependent long-range facilitation and suppression in the human visual cortex. Vision Res. 36, 2099–2109 (1996)

13. Polat, U., Sagi, D.: The architecture of perceptual spatial interactions. Vision Res. 34, 73–78 (1994)

14. Angelucci, A., Bressloff, P.C.: Contribution of feedforward, lateral and feedback connections to the classical receptive field center and extra-classical receptive field surround of primate V1 neurons. Prog. Brain Res. 154, 93–120 (2006)

15. Angelucci, A., Bijanzadeh, M., Nurminen, L., Federer, F., Merlin, S., Bressloff, P.C.: Circuits and mechanisms for surround modulation in visual cortex. Annu. Rev. Neurosci. 40, 425–451 (2017)

16. Gilbert, C.D.: Horizontal integration in the neocortex. Trends Neurosci. 8, 160–165 (1985)

17. Gilbert, C.D., Wiesel, T.N.: Morphology and intracortical projections of functionally characterised neurones in the cat visual cortex. Nature 280, 120–125 (1979)

18. Rockland, K.S., Lund, J.S.: Intrinsic laminar lattice connections in primate visual cortex. J. Comp. Neurol. 216, 303–318 (1983)

19. Livingstone, M.S., Hubel, D.H.: Specificity of intrinsic connections in primate primary visual cortex. J. Neurosci. 4, 2830–2835 (1984)

20. Martin, K., Whitteridge, D.: Form, function and intracortical projections of spiny neurones in the striate visual cortex of the cat. The Journal of physiology 353, 463–504 (1984)

21. Palmer, L.A., Davis, T.L.: Receptive-field structure in cat striate cortex. J. Neurophysiol. 46, 260–276 (1981)

22. Adams, D.L., Horton, J.C.: Ocular dominance columns in strabismus. Vis. Neurosci. 23, 795–805 (2006)

23. Shatz, C.J., Stryker, M.P.: Ocular dominance in layer IV of the cat’s visual cortex and the effects of monocular deprivation. The Journal of Physiology 281, 267–283 (1978)

24. Lowel, S.: Ocular dominance column development: strabismus changes the spacing of adjacent columns in cat visual cortex. J. Neurosci. 14, 7451–7468 (1994)

25. Tso, D. Y., Gilbert, C. N W.T.,: Relationships between horizontal interactions and functional architecture in cat striate cortex as revealed by cross-correlation analysis. J. Neurosci. 3, 1160–1170 (1986)

26. Ts’o, D., Gilbert, C.D.: The organization of chromatic and spatial interactions in the primate striate cortex. J. Neurosci. 8, 1712–1727 (1988)

27. Gilbert, C.D., Wiesel, T.N.: Columnar specificity of intrinsic horizontal and corticocortical connections in cat visual cortex. Journal of Neuroscience 9, 2432–2442 (1989)

28. Gilbert, C.D., Das, A., Ito, M., Kapadia, M., Westheimer, G.: Spatial integration and cortical dynamics. Proceedings of the National Academy of Sciences 93, 615–622 (1996)

29. Brainard, D.H.: The Psychophysics Toolbox. Spat. Vis. 10, 433–436 (1997)

30. Pelli, D.G.: The VideoToolbox software for visual psychophysics: transforming numbers into movies. Spat. Vis. 10, 437–442 (1997)

31. Sereno, M.I., Dale, A.M., Reppas, J.B., Kwong, K.K., Belliveau, J.W., Brady, T.J., Rosen, B.R., Tootell, R.B.: Borders of multiple visual areas in humans revealed by functional magnetic resonance imaging. Science 268, 889–893 (1995)

32. Polimeni, J.R., Bhat, H., Witzel, T., Benner, T., Feiweier, T., Inati, S.J., Renvall, V., Heberlein, K., Wald, L.L.: Reducing sensitivity losses due to respiration and motion in accelerated echo planar imaging by reordering the autocalibration data acquisition. Magn. Reson. Med. (2015)

33. Fischl, B.: FreeSurfer. Neuroimage 62, 774–781 (2012)

34. Wang, J., Nasr, S., Roe, A.W., Polimeni, J.R.: Critical factors in achieving fine-scale functional MRI: Removing sources of inadvertent spatial smoothing. Wiley Online Library (2022)

35. Greve, D.N., Fischl, B.: Accurate and robust brain image alignment using boundary-based registration. Neuroimage 48, 63–72 (2009)

36. Blazejewska, A.I., Fischl, B., Wald, L.L., Polimeni, J.R.: Intracortical smoothing of small-voxel fMRI data can provide increased detection power without spatial resolution losses compared to conventional large-voxel fMRI data. Neuroimage 189, 601–614 (2019)

37. Fracasso, A., Dumoulin, S.O., Petridou, N.: Point-spread function of the BOLD response across columns and cortical depth in human extra-striate cortex. Prog. Neurobiol. 207, 102187 (2021)

38. Olman, C., Van de Moortele, P., Ugurbil, K.: Point spread function for gradient echo and spin echo BOLD fMRI at 7 Tesla. In: ISMRM 12th Scientific Meeting and Exhibition, Kyoto, Japan, pp. 1066. (2004)

39. Adams, D.L., Sincich, L.C., Horton, J.C.: Complete pattern of ocular dominance columns in human primary visual cortex. J. Neurosci. 27, 10391–10403 (2007)

40. Tootell, R.B., Hamilton, S.L., Silverman, M.S., Switkes, E.: Functional anatomy of macaque striate cortex. I. Ocular dominance, binocular interactions, and baseline conditions. J. Neurosci. 8, 1500–1530 (1988)

41. Sincich, L.C., Adams, D.L., Horton, J.C.: Complete flatmounting of the macaque cerebral cortex. Vis. Neurosci. 20, 663–686 (2003)

42. Hubel, D.H., Wiesel, T.N., LeVay, S.: Functional architecture of area 17 in normal and monocularly deprived macaque monkeys. Cold Spring Harb. Symp. Quant. Biol. 40, 581–589 (1976)

